# TCF4 Mediates PHF6 Transcriptional Regulation of Neural Stem Cells in the Developing Brain, a Mechanism Disrupted in Börjeson–Forssman–Lehmann Syndrome (BFLS)

**DOI:** 10.64898/2026.09.12.750495

**Authors:** Behnaz Nateghi, Yunqiao Li, Dianbo Qu, Dilan Rasool, Ariel Madrigal, Hamed S Najafabadi, Pierre Mattar, Vahab D. Soleimani, Azad Bonni, Arezu Jahani-Asl

**Author notes:** These authors contributed equally to this work (Behnaz Nateghi, Yunqiao Li).

## Abstract

Transcriptional hierarchies govern neural stem cell (NSC) fate decisions during neurodevelopment, and disruption of these regulatory networks underlies neurodevelopmental disorders. Börjeson-Forssman-Lehmann syndrome (BFLS) is an X-linked intellectual disability (XLID) caused by mutations in a chromatin-associated transcriptional regulator, the plant homeodomain zinc finger protein 6 (*PHF6*). Although BFLS mouse models harboring *PHF6* patient mutations recapitulate neurogenic and cognitive deficits, the molecular mechanisms linking PHF6 dysfunction to these phenotypes remain poorly understood. Here, we identify transcription factor 4 (TCF4), a basic helix–loop–helix transcription factor implicated in neurodevelopmental and psychiatric disorders, as a critical downstream effector of PHF6 in NSC fate regulation. *Tcf4* expression is reduced in BFLS mouse models harbouring R342X and C99F- m mutations and in a conditional *Phf6* knockout driven by Nestin-Cre. Mechanistically, PHF6 binds the *Tcf4* gene regulatory element upstream of its transcriptional start site (TSS) and promotes *Tcf4* expression. *Tcf4* depletion in embryonic NSCs enhances self-renewal and stemness, increasing neurosphere formation and expression of stem cell markers, *Nestin* and *Sox2*. Conversely, TCF4 restoration rescues NSC defects caused by PHF6 loss-of-function in BFLS and *Phf6*/Nestin-Cre mice. This regulatory relationship persists in adult NSCs where TCF4 similarly restricts stem cell expansion. Importantly, restoration of TCF4 expression in embryonic brain ameliorates behavioral and cognitive deficits in BFLS mice. Together, these findings establish a PHF6-*Tcf4* transcriptional pathway that restricts NSC self-renewal and links disrupted transcriptional control of NSC fate to the neurodevelopmental and behavioral abnormalities of BFLS.

## Introduction

X-linked intellectual disability (XLID) affects approximately 1-3% of the population^1–4^ and is clinically characterized by a deficit in intellectual function with an intelligence quotient (IQ) below 70 before 18 years of age^5, 6^. Genetic aberrations on the X chromosome, such as gene deletions, duplications, inversions, and point mutations, have been associated with XLID^7, 8^. Börjeson-Forssman-Lehmann syndrome (BFLS) is an XLID caused by mutations in the plant homeodomain zinc finger protein 6 (*PHF6*)^9–12^. BFLS patients present with intellectual disabilities, ranging from mild to severe, with seizure susceptibility^9, 11, 13^. Additional physical phenotypic characteristics include short stature, broad forehead, deep-set eyes, and obesity^9, 11, 14^. Multiple mutations in the *PHF6* gene on the X chromosome have been identified in BFLS patients^11, 13, 15, 16^, with five recurrent mutations reported, including R342X, accounting for 21% of cases^10, 17, 18^.

*Phf6* gene is located at Xq26-27 in humans and is composed of 11 exons^10, 11, 19^. Structurally, *PHF6* contains two PHD zinc-like finger domains and consists of two nuclear and one nucleolar localization sequence^10, 20^. *PHF6* is highly conserved in vertebrates, with high expression during early fetal life and embryonic tissue specifically in the developing central nervous system during the early stages of corticogenesis^21, 22^. Mutations in the evolutionarily conserved *PHF6* cysteine residues, C45, C99, and C305, in either the C2HC or PHD-type zinc finger, impair accurate protein folding, leading to PHF6 loss of function as reported in the BFLS studies^10, 20^.

Transcription factor 4 (TCF4), also known as ITF2 or E2-2, belongs to the family of basic Helix-Loop-Helix (bHLH) transcription factors (TFs)^23, 24^. Similar to PHF6, TCF4 plays critical roles in a wide range of neurodevelopmental processes, including neural precursor proliferation, differentiation, and migration^25–28^. TCF4 is abundantly expressed across different regions of the brain and exhibits layer-specific expression patterns in the developing murine neocortex, suggesting a pivotal role in orchestrating brain development^28^. In the adult brain, TCF4 is highly expressed in the hippocampal dentate gyrus, a region where neural stem/progenitor cells generate new functional neurons throughout life^29, 30^. Emerging evidence is mounting that TCF4 deregulation is associated with neurodevelopmental and psychiatric disorders. For example, heterozygous loss-of-function mutations in *TCF4* cause Pitt-Hopkins syndrome (PTHS)^30–32^, and genome-wide association studies (GWAS) have identified *TCF4* polymorphisms associated with schizophrenia and other psychiatric conditions^33–35^.

In this study, we report a novel PHF6-*Tcf4* signalling axis in the regulation of NSCs, a mechanism perturbed in BFLS. Via the intersection of public PHF6 chromatin immunoprecipitation sequencing (ChIP-Seq)^36^ and RNA-Seq followed by GO Term functional annotation, we identified TCF4 as a high-confidence direct candidate target gene of PHF6. Using genetic mouse models, including BFLS patient mouse models and *Phf6* conditional knockout (KO) model, we consistently found significant downregulation of TCF4 mRNA and protein expression. Functional assays revealed that, similar to PHF6, TCF4 restricts NSCs self-renewal in the developing brain, a mechanism impaired in BFLS. Importantly, we demonstrate that PHF6 directly binds to the *Tcf4* promoter to upregulate its expression in NSCs, and forced expression of TCF4 rescues the PHF6-loss-of-function defects in BFLS-derived NSCs. The behavioral characterization of BFLS mouse models showed a consistent behavioral phenotype characterized by hyperlocomotor activity and cognitive impairments, including deficits in object novelty recognition. Notably, in vivo restoration of TCF4 expression through in utero electroporation (IUE) significantly improved these behavioral and cognitive abnormalities, providing functional evidence linking dysregulation of the PHF6–*Tcf4* molecular pathway to the behavioral phenotypes observed in BFLS mouse models. Our data suggest that impairment of the PHF6-*Tcf4* pathway underlies defects in neurogenesis. These findings not only provide the first line of evidence on TCF4 deregulation in BFLS, but also open up new avenues of investigation to assess PHF6 regulation of neurogenesis in a broad range of psychiatric and neurodevelopmental disorders.

## Results

### Gene expression analyses of PHF6 and TCF4 during cortical neurogenesis

In analysis of PHF6 genome-wide occupancy using ChIP-Seq, we previously established that PHF6 directly binds DNA in the embryonic mouse brain^36^. Via mapping PHF6-differentially regulated genes in cortical progenitors and its intersection with PHF6-bound loci, we demonstrated that PHF6 operates as a regulator of neurogenesis ^36^. Building on these findings, we sought to map PHF6 mechanisms, focusing on downstream effectors that mediate PHF6 regulation of NSC fate. This investigation revealed robust binding of PHF6 at the *Tcf4* promoter, a TF that is implicated in neurodevelopmental disorders (p = 7.32 × 10^⁻536^). Consistent with this observation, reanalysis of published RNA-seq datasets demonstrated reduced *T*cf4 expression following *Phf6* knockdown^36^, supporting that *Tcf4* may serve as a candidate downstream target of PHF6. To further characterize the relationship between *Phf6* and *Tcf4* during neurodevelopment, we examined a publicly available single-cell RNA-seq atlas of the developing mouse cortex^37^. We found that both genes were broadly expressed across neurodevelopmental stages, spanning embryonic day 10 (E10) to postnatal day 4 (P4) (**Fig. 1A-C**). Pearson correlation analysis of imputed gene expression values demonstrated a positive association between *Phf6* and *Tcf4* across multiple cortical cell populations, including apical progenitors and cycling glial cells (**Fig. 1D, E**). To determine whether *Phf6* and *Tcf4* exhibit coordinated temporal expression across developmental time, we also examined normalized expression across progenitor populations. Both genes showed sustained expression across embryonic stages, with enrichment in apical progenitors, intermediate progenitors, and cycling glial cells (**Fig. 1F, G**). Overall, *Phf6* and *Tcf4* exhibited similar developmental trajectories across multiple lineages, spanning from early neurogenesis through late embryonic stages, raising the question of whether these two transcription factors, known to be associated with different human disease, operate on the same pathway to regulate neurogenic processes.

**Figure 1:**
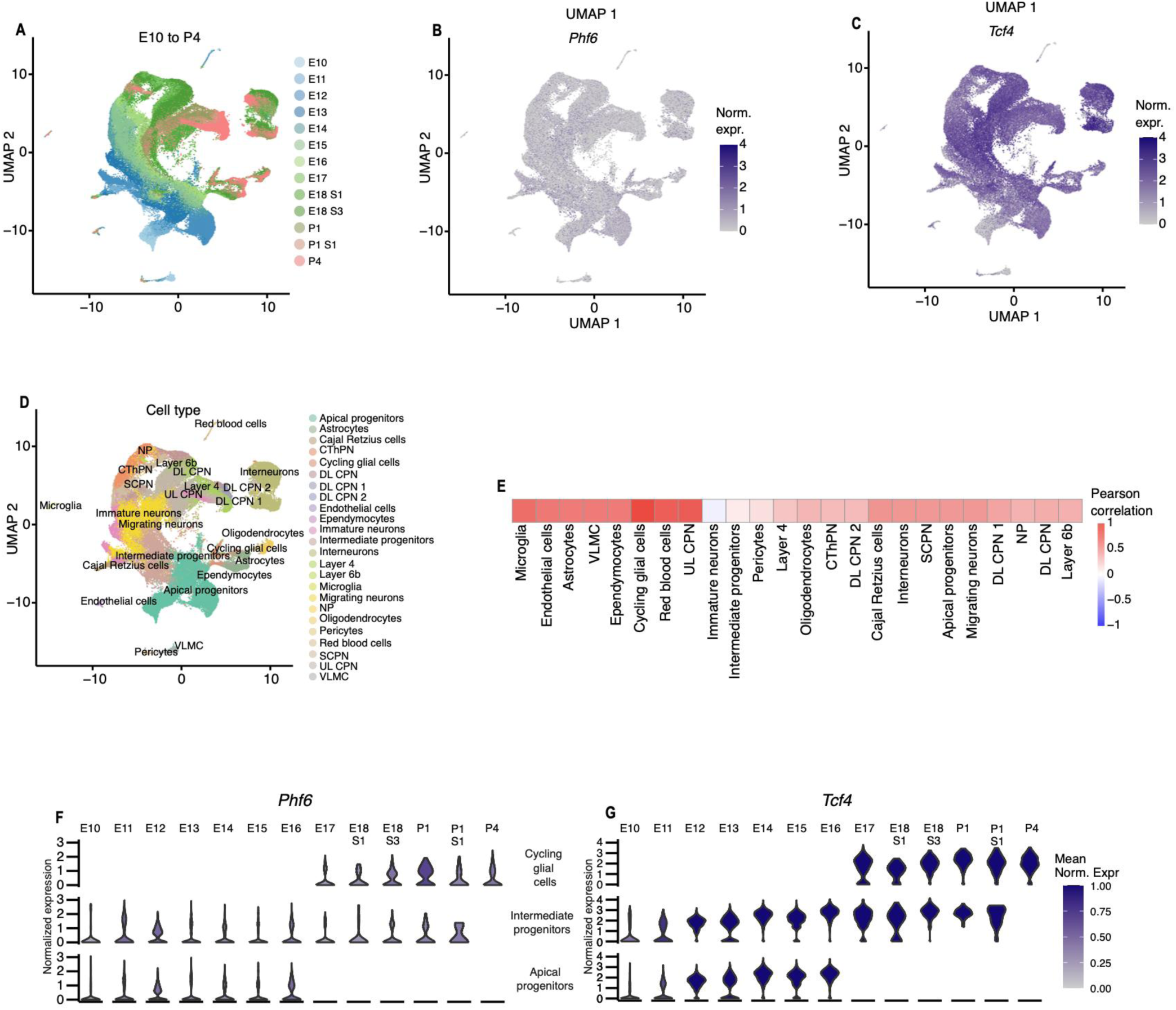
Cell type-specific co-expression analysis of *Phf6* and *Tcf4* in mouse cerebral cortex. Low-dimensional representation of single cells from the mouse cerebral cortex, derived from UMAP embedding of single-cell RNA-seq data^37^ are shown. (A) animal age, or expression level of (B) *Phf6* and (C) *Tcf4*. (D) cell-typed annotations. (E) Heatmap displaying Pearson correlation coefficients between *Phf6* and *Tcf4* across various cell types. (F-G) Violin plots showing the expression of *Phf6* (F) and *Tcf4* (G) in the developing mouse cerebral cortex. Violin color represents the mean normalized expression level for each cell type at each developmental stage. Correlation values were computed using imputed gene expression profiles obtained after applying MAGIC^55^. Single cell RNA-seq data was obtained from Di Bella et al, 2021^37^.

### TCF4 expression is attenuated upon deletion of *Phf6* in Nestin-expressing cells

We set out to characterize how TCF4 expression is altered in eNSCs lacking PHF6. We induced genetic deletion of *Phf6* via breeding *Phf6*^loxP/loxP^ with Nestin-CreERT2^+^ mice, and tamoxifen administration at E12 to delete *Phf6* exons 4 and 5 in Nestin-expressing cells embryonically, followed by analysis of brain sections at E14 (**Fig. 2A**). Immunohistochemical analysis of the embryonic brain sections revealed consistent expression of TCF4 in the subventricular zone (SVZ) of Phf6^loxP/Y^/Nestin-CreERT2^−^ control (**Fig. 2B**), consistent with its role in late embryonic and early postnatal neurogenesis^28, 38^. In contrast, TCF4 expression was markedly reduced upon deletion of *Phf6*, as revealed by analysis of brain sections from the in Phf6^-/Y^/Nestin-CreERT2^+^ mice (**Fig. 2B**). To validate our observation, we next examined the mRNA and protein expression levels of TCF4 in brain tissue. RT-qPCR analysis revealed a significant reduction in *Tcf4* mRNA expression at E14 in Phf6^-/Y^/Nestin-CreERT2^+^ mice compared to the Phf6^loxP/Y^/Nestin-CreERT2^−^ control (**Fig. 2C**). Consistent with these data, immunoblotting of E14 cortical lysates of Phf6^-/Y^/Nestin-CreERT2^+^ model revealed a marked reduction in TCF4 protein levels (**Fig. 2D, E**). Together, our data revealed that PHF6 deletion in Nestin-expressing cells results in a significant reduction in TCF4 expression.

**Figure 2:**
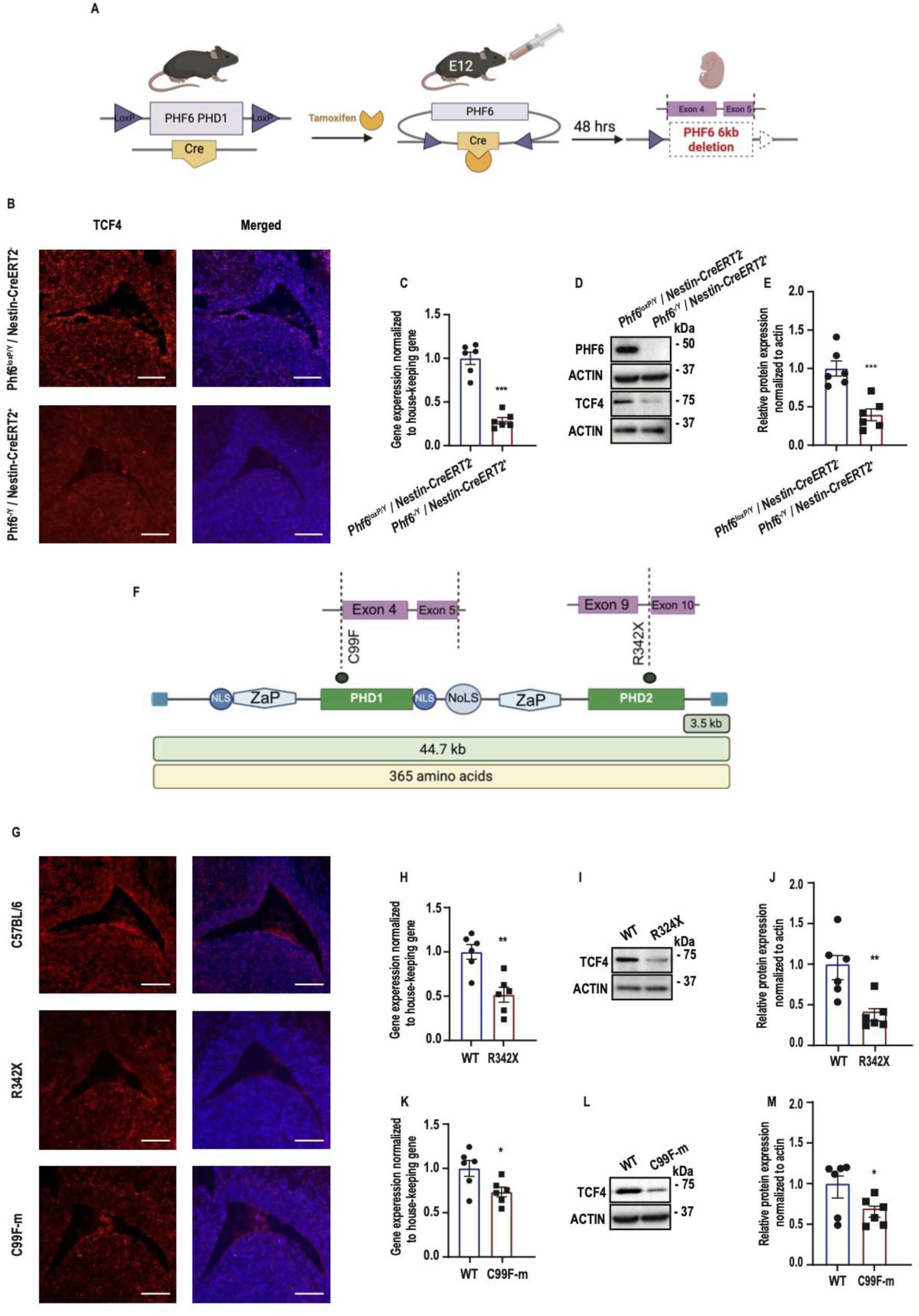
TCF4 expression is impaired in BFLS mice. (A) A floxed allele of PHF6, with LoxP sites flanking the PHD1 domain, was used to generate a conditional knockout mouse model. Upon tamoxifen administration at E14, Cre recombinase driven by the Nestin-CreERT2 promoter is activated and translocates to the nucleus, where it mediates site-specific recombination at the LoxP sites. This results in a 6 kb genomic deletion of the PHF6 gene at exons 4 and 5. Made with BioRender. (B) Immunofluorescence (IF) staining of coronal sections from E14 for Phf6^-/Y^/Nestin-CreERT2^+^ and control Phf6^loxP/Y^/Nestin-CreERT2^-^ male mice using a TCF4 antibody (red). (C-E) mRNA and protein of Phf6^-/Y^/Nestin-CreERT2^+^ and control Phf6^loxP/Y^/Nestin-CreERT2^-^ mice ∼E14 were subjected to RT-qPCR and immunoblotting analysis (n=3). (F) Illustration of the full *Phf6* gene. The two recurrent BFLS mutations are shown: C99F and R342X (truncation at exon 10). Made with BioRender. (G) Immunofluorescence (IF) staining of coronal sections from E14 for R342X, C99F-m, and littermate C57BL/6 wild-type (WT) males as controls using a TCF4 antibody (red). (H-M) mRNA and protein of R342X, C99F-m, and wild-type control mice ∼E14 were subjected to RT-qPCR and immunoblotting analysis (n=3). Statistical comparisons were performed using an unpaired two-tailed Student’s t-test. Data information: Data are presented as mean ± SEM. *p < 0.05, **p < 0.01, ***p < 0.001. n represents an independent biological sample.

### TCF4 expression is reduced in BFLS mouse models

To address whether PHF6 regulation of TCF4 expression is impaired in the disease context, we set out to assess TCF4 expression in two BFLS mouse models harboring *PHF6* patient mutations, R342X and C99F-m. The R342X mutation is the most recurrent BFLS patient-mutation occurring at exon 10 (C.1024 C > T), impairing the ePHD2 domain, whereby PHF6 is proposed to function as a truncated protein^9–12, 17, 18, 39, 40^. Another BFLS patient point mutation (m) in *Phf6* is wherein cysteine-99 is replaced with phenylalanine (C99F), caused by a nucleotide change at position 296 (G>T), impairing the function of the PHD1 domain (**Fig. 2F**). We subjected brain sections from R342X and C99F-m to immunostaining analysis. Similar to results obtained in PHF6 cKO model, we observed a reduction in TCF4 expression at E14 in both R342X and C99F-m mutant models, with the most pronounced effects in the brain of R342X mice (**Fig. 2G**). RT-qPCR analysis revealed a significant decrease in *Tcf4* mRNA in both R342X and C99F-m brain tissue compared to control mice (**Fig. 2H, I**). Moreover, immunoblotting analysis showed that both R342X and C99F-m mutants displayed reduced TCF4 protein expression compared to corresponding controls (**Fig. 2J, K-M**). Together, our data using three different mouse models reveal that genetic deletion of *Phf6* or its loss-of-function mutations impairs the expression of TCF4, raising the question of whether TCF4 is a downstream effector of PHF6 in the regulation of neurogenesis.

### TCF4 is a direct transcriptional target of PHF6

Having established the significant deregulation of TCF4 expression in a PHF6-dependent manner in BFLS and Phf6^-/Y^/Nestin-CreERT2^+^ mice, we next asked if TCF4 is a direct transcriptional target of PHF6. We performed ChIP-qPCR in embryonic cortical tissue, utilizing an antibody to endogenous PHF6 for chromatin precipitation. A non-specific IgG antibody was used in parallel as a negative control to assess the background signal. Following immunoprecipitation, we performed qPCR using primers designed against the *Tcf4* peak summit at the promoter, in which PHF6 binding was predicted from prior ChIP-seq analysis^36^. As controls, we included EphA4, a known PHF6 target^36^, as a positive control, and Zfp735^36^, a genomic locus with no predicted PHF6 occupancy, as a negative control. In cortices from wild-type mice, PHF6 was significantly enriched at *Tcf4* loci (**Fig. 3A**), supporting robust PHF6 binding to the *Tcf4* promoter. To assess whether this regulatory interaction is disrupted in BFLS models, we performed ChIP-qPCR on E14 cortical tissues from Phf6^-/Y^/Nestin-CreERT2^+^, R342X, and C99F-m mice. In Phf6^-/Y^/Nestin-CreERT2^+^ mice, PHF6 binding at the *Tcf4* promoter was markedly reduced compared to control littermates (**Fig. 3B**). Similarly, PHF6 occupancy at *Tcf4* promoters significantly decreased in R342X (**Fig. 3C**) and C99F-m (**Fig. 3D**) cortices, consistent with the loss of DNA-binding ability in these mutant mice. Our findings demonstrate that PHF6 directly binds to the *Tcf4* promoter *in vivo* and this interaction is consistently disrupted across different mouse models upon PHF6 loss-of-function.

**Figure 3:**
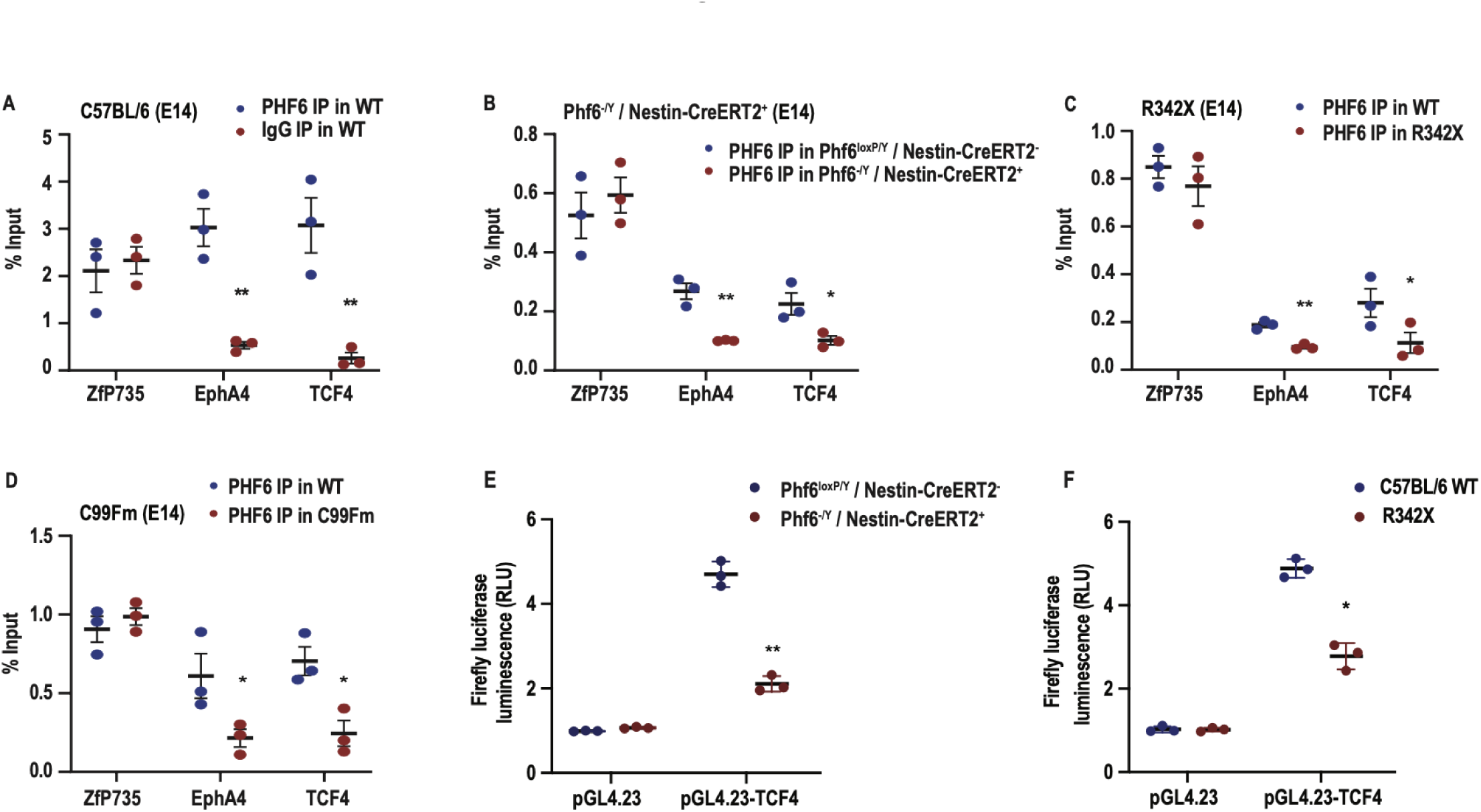
TCF4 is a direct transcriptional target of PHF6. (A-D) Cerebral cortical tissues were isolated from Phf6^-/Y^ / Nestin-CreERT2^+^ (cKO), Phf6^loxP/Y^ / Nestin-CreERT2^-^ (control); R342X, and C99F-m and their littermate C57BL/6 as controls at E14. Samples were subjected to ChIP-PCR using a PHF6 antibody. Zfp735 loci were used as a negative control, and EphA4 as a positive control for the PCR. (E-F) Dual luciferase reporter assay was performed in eNSC 48 h following electroporation with a firefly luciferase reporter plasmid containing the TCF4 promoter or the control pGL4.23-basic vector. Statistical comparisons were performed using an unpaired two-tailed Student’s t-test. Data information: Data are presented as mean ±SEM. *p < 0.05, **p < 0.01, ***p < 0.001. Representative data of n=3 independent replicates are shown in panels. n represents an independent biological sample.

To assess whether the direct binding of PHF6 to *Tcf4* promoter induces *Tcf4* expression, we designed a dual-luciferase reporter assay. PHF6 sequence within the peak identified by ChIP-seq^36^ were cloned into the pGL4.23 luciferase reporter vector, and the constructs were verified by DNA sequencing. We performed a dual-luciferase reporter assay in eNSCs from Phf6^-/Y^/Nestin-CreERT2^+^ and R342X mutant mice. Cells were electroplated with a firefly luciferase reporter plasmid containing the *Tcf4* promoter or the control pGL4.23-basic vector, and a Renilla luciferase expression plasmid. Luciferase activity was evaluated 48 hours post-electroporation. Our results demonstrated that PHF6 regulation of *Tcf4* transcriptional activity was significantly impaired in the Phf6^-/Y^/Nestin-CreERT2^+^ compared to Phf6^loxP/Y^/Nestin-CreERT2^−^ control (**Fig. 3E**). Similarly, eNSCs from the R342X exhibited a significant reduction in the luciferase activity (**Fig. 3F**), supporting that PHF6 directly upregulates *Tcf4* transcription, and PHF6 genetic deletion or truncation significantly attenuates *Tcf4* expression.

### TCF4 restricts eNSC self-renewal and rescues PHF6-deficient-eNSCs phenotype

PHF6 restricts eNSC self-renewal, and its loss of function promotes eNSCs expansion^36, 41^, suggesting its involvement in the regulation of NSCs and lineage commitment. Having shown that PHF6 directly binds *Tcf4* promoter to upregulate its expression, led us next to the question of whether genetic knockdown (KD) of *Tcf4* functionally phenocopies PHF6-loss-of-function defects in the regulation of NSCs.

We employed an siRNA-mediated approach in primary E14 wild-type (WT)- eNSCs and confirmed efficient KD of *Tcf4* (**Fig. 4A**). To evaluate NSC self-renewal, we conducted an extreme limiting dilution assay (ELDA)^42, 43^ to assess the self-renewal capacity of eNSCs. Our results revealed a significant increase in eNSC self-renewal upon knockdown of *Tcf4* (**Fig. 4B, C**). In parallel, we also conducted a limiting dilution assay (LDA), which demonstrated an increase in the number of neurosphere formation in *Tcf4* KD compared to non-targeting control (**Fig. 4D**). Consistent with these data, sphere size analysis revealed a significant increase in diameter in *Tcf4* KD cultures (133.1 μm) compared to controls (61.1 μm) (**Fig. 4E**), further supporting a role of TCF4 in restricting eNSC expansion. Next, we subjected the *Tcf4* KD and control eNSCs to RT-qPCR analysis to evaluate changes in the mRNA expression of stem cell markers, including *Sox2* and *Nestin*. We found that the KD of *Tcf4* led to a significant increase in the mRNA levels of both genes (**Fig. 4F**). To determine whether these changes in gene expression were reflected at the protein level, we conducted immunoblotting analysis, and detected elevated protein levels of SOX2 and NESTIN in *Tcf4* KD cells compared to non-targeting control siRNA (**Fig. 4G, H**). Collectively, these findings demonstrate that *Tcf4* KD enhances eNSC self-renewal and stemness, mirroring the effects of PHF6 loss. This supports a model in which *Tcf4* may operate downstream of PHF6 to regulate NSC fate during embryonic brain development. To assess this model, we asked whether forced expression of TCF4 rescues the NSC defects observed in PHF6-deficient cultures. We conducted electroporation of TCF4 or a vector control in eNSC obtained from *Phf6^-/Y^/Nestin- CreERT2^+^*, and subjected the cultures to a series of functional and molecular assays to assess self-renewal and stemness. ELDA revealed that forced expression of TCF4 significantly suppressed aberrant self-renewal in *Phf6^-/Y^/Nestin-CreERT2^+^* eNSCs (**Fig. 4I**). Consistently, LDA showed a decrease in neurosphere-forming frequency in TCF4-expressing cells compared to controls (**Fig. 4J**). In parallel, we examined the mRNA expression of *Sox2* and Nestin, and found significant attenuation at transcript levels for both markers in TCF4-expressing cells relative to control (**Fig. 4K**). This was further supported by immunoblotting, which confirmed reduced protein levels of SOX2 and NESTIN (**Fig. 4L**), indicating that TCF4 forced expression suppresses the expression of stemness markers.

**Figure 4:**
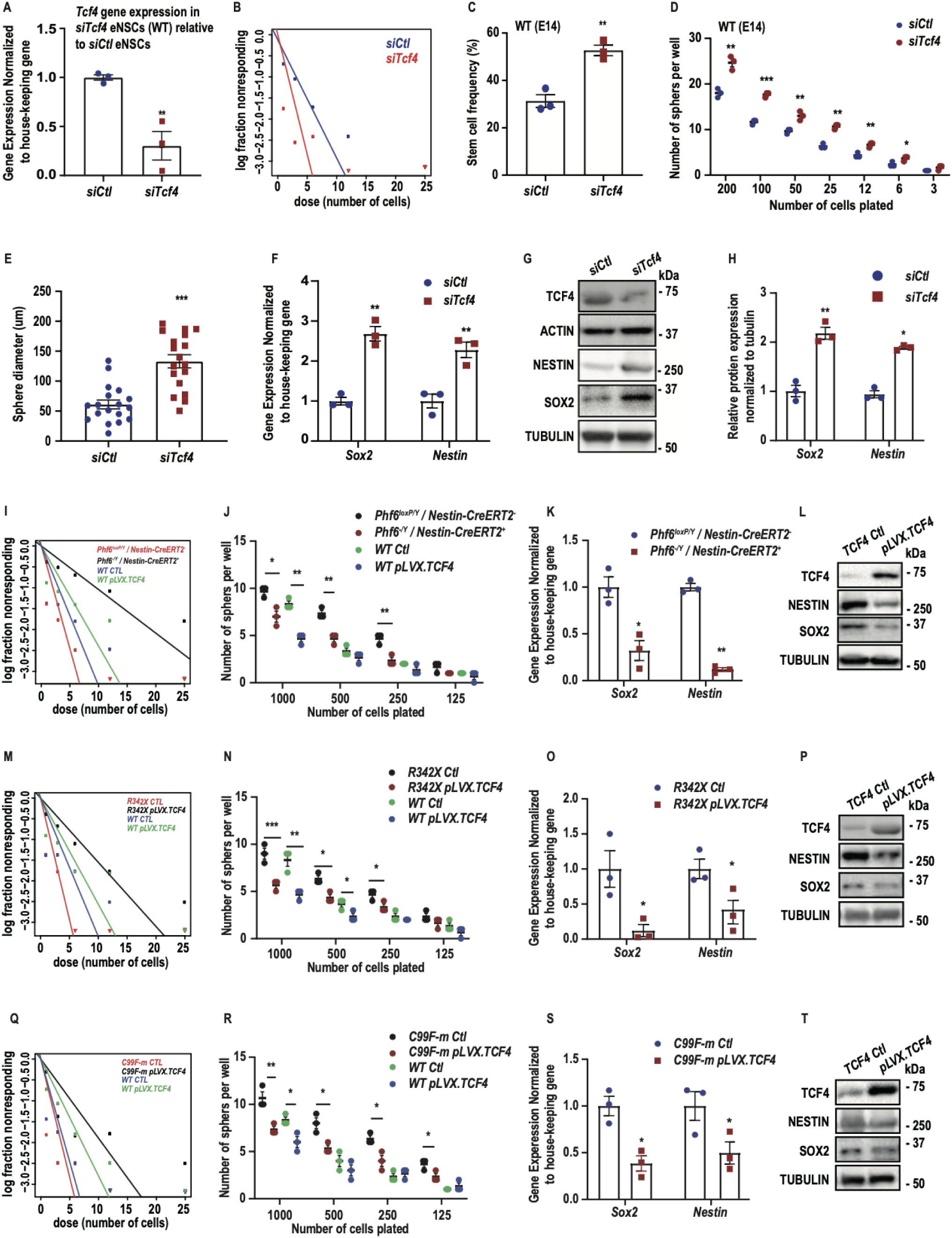
Knockdown of *Tcf4* impairs NSCs in the embryonic brain, and forced expression of TCF4 rescues these defects. eNSCs were isolated and cultured from WT mice at E14, and *Tcf4* KD was induced using an siRNA approach (A). (B-D) Samples were analyzed using an extreme limiting dilution assay (ELDA) and limiting dilution assay (LDA) 7 days post-plating. (E) sphere diameter, (F) RT-qPCR analysis using Nestin and Sox2 primers. (G, H) Immunoblotting using NESTIN and SOX2 antibodies was also performed 7-days post-plating. Phf6 cKO, R342X, C99F-m, and WT eNSCs cultured at E14 were electroporated with pLVX.TCF4 and samples were subjected to (I, M, Q) ELDA, (G, N, R) LDA, (K, O, S) RT-qPCR, and (L, P, T) immunoblotting analysis with SOX2 and NESTIN. Statistical analyses were performed using unpaired two-tailed Student’s t-test or one-way ANOVA, where appropriate. Data information: Data are presented as mean ±SEM. *p < 0.05, **p < 0.01, ***p < 0.001. Representative plots of n=3 independent replicates.

### TCF4 reverses the BFLS-derived eNSCs phenotype

Next, we assessed whether TCF4 rescues PHF6-deficient phenotypes in the R342X model. We induced the expression of TCF4 in eNSCs derived from R342X mutant mice. TCF4 forced expression led to a significant reduction in neurosphere formation, as assessed by ELDA and LDA (**Fig. 4M, N**). Consistently, expression levels of the stemness markers SOX2 and NESTIN were significantly attenuated at both mRNA and protein levels upon induction of TCF4 (**Fig. 4O, P**). Similarly, in examining the effect of TCF4 in eNSCs derived from C99F-m mutant mice, enforced expression of TCF4 significantly reduced self-renewal and neurosphere formation, as assessed by ELDA and LDA, respectively (**Fig. 4Q, R**). In addition, TCF4 forced expression resulted in a marked downregulation of SOX2 and NESTIN at both the transcript and protein levels (**Fig. 4S, T**). Together, these findings demonstrate that TCF4 can mitigate the aberrant self-renewal phenotype caused by PHF6 deficiency or loss of function mutations. Our data suggests that TCF4 is a key functional downstream effector of PHF6 in regulating eNSC fate.

### TCF4 regulates stemness in adult BFLS-derived NSCs

Given the established role of TCF4 in NSC regulation^44^ and its high expression in adult neurogenic niches^30, 45^, we next investigated whether its expression is altered in adult BFLS mouse models. RT-qPCR analysis of adult NSCs (aNSCs) derived from R342X and C99F-m revealed that *Tcf4* transcript levels were significantly reduced in both models compared to corresponding wild-type controls (**Fig. 5A**), with the R342X line showing a more pronounced reduction (p = 0.01).

**Figure 5:**
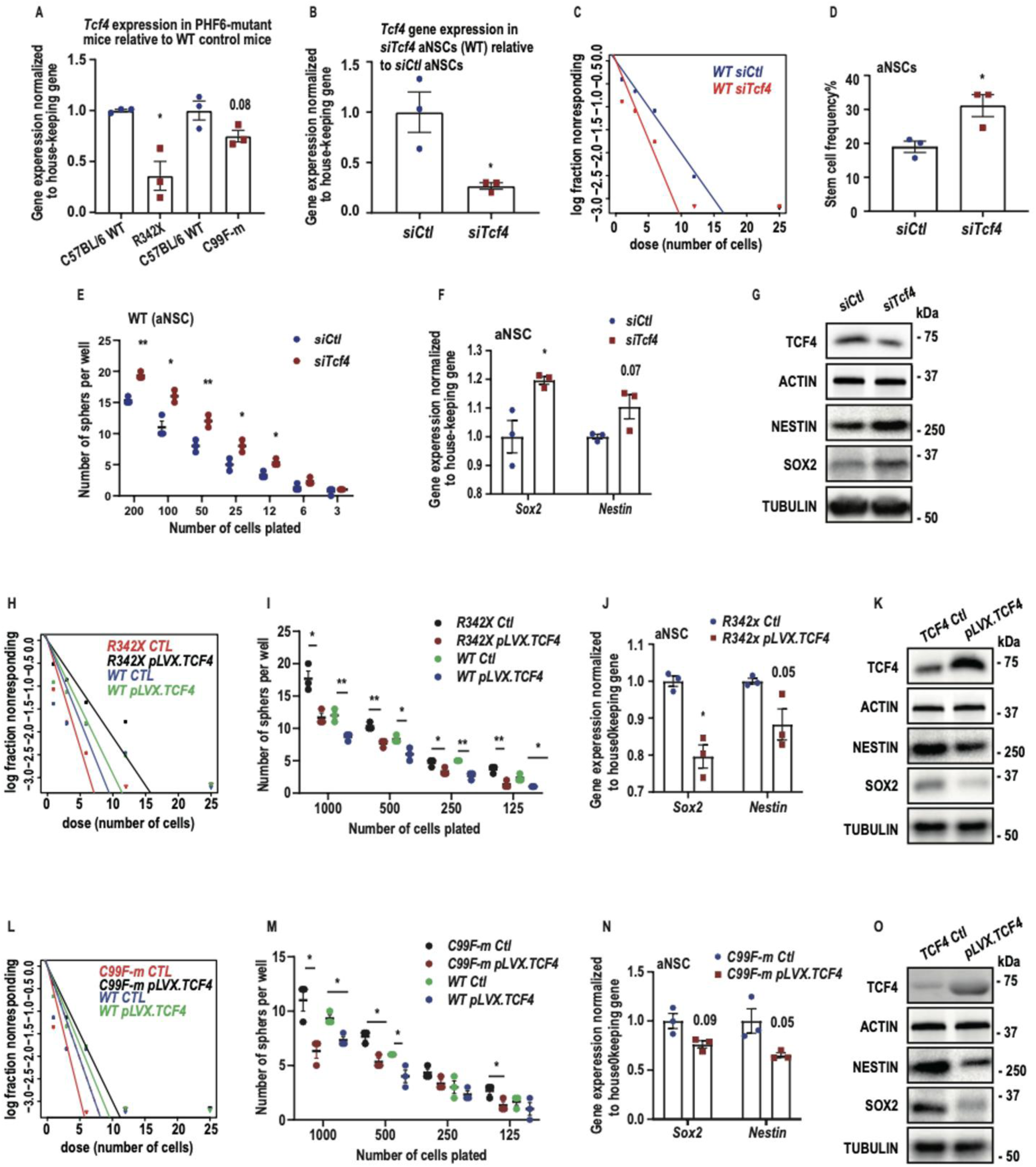
TCF4 regulates stemness in adult BFLS-derived NSCs. (A) RT-qPCR was performed to investigate the expression level of *Tcf4* in PHF6 mutant mouse models. (B) The aNSCs were isolated and cultured from WT mice and Tcf4 KD was induced using a siRNA approach. (C-E) Samples were analyzed using ELDA and LDA. (F) RT-qPCR analysis using primers targeting Sox2 and Nestin, (G), and immunoblotting using NESTIN and SOX2 antibodies. R342X, C99F-m, and WT aNSCs cultured were electroporated with pLVX.TCF4 and samples were subjected to (H, L) ELDA, (I, M) LDA. (G, N) stem cell markers were examined using RT-qPCR using Sox2 and Nestin primers, and (K, O) immunoblotting analysis with SOX2 and NESTIN antibodies. Statistical analyses were performed using unpaired two-tailed Student’s t-test or one-way ANOVA, where appropriate. Data information: Data are presented as mean ± SEM. *p < 0.05, **p < 0.01, ***p < 0.001.

To examine the functional consequences of *Tcf4* deficiency, we performed siRNA-mediated KD studies in aNSCs derived from wild-type mice (**Fig. 5B**). ELDA revealed that *Tcf4* KD significantly increased self-renewal and stem cell frequency (**Fig. 5C, D**). Similarly, LDA showed an increased number of neurospheres across multiple dilutions in *Tcf4*-deficient aNSCs (**Fig. 5E**), reinforcing the conclusion that *Tcf4* loss promotes aNSC expansion. To uncover the molecular basis of this phenotype, we analyzed *Sox2* and *Nestin* expression using RT-qPCR and immunoblotting. Both mRNA and protein levels of these stemness markers were elevated in *Tcf4* KD aNSCs compared to controls (**Fig. 5F, G**), supporting that TCF4 negatively regulates stemness expansion. Overall, these findings indicate that TCF4 downregulation in aNSCs contributes to increased self-renewal, highlighting a key role for TCF4 in regulation of aNSCs behavior.

Building on our observation that *Tcf4* expression is reduced in PHF6 mutant aNSCs, we next asked whether TCF4 forced expression could reverse the phenotype of aNSCs from R342X and C99F-m mice. aNSCs electroporated with TCF4 or empty vector control were subjected to ELDA, LDA, and analysis of stem cell markers. Similar to our results obtained from eNSCs, forced expression of TCF4 resulted in a marked decrease in neurosphere-forming frequency (**Fig. 5H, I**), and a marked reduction in the expression of SOX2 and NESTIN (**Fig. 5J, K**). Similarly, TCF4 rescued aNSCs defects derived from C99F-m mice (**Fig. 5**L-**O**). These data demonstrate that *Tcf4* KD significantly increases NSC self-renewal, indicating a requirement for TCF4 in the maintenance of normal NSC commitment levels.

### TCF4 restoration rescues altered locomotor activity in BFLS

To examine behavioral abnormalities associated with PHF6 mutations and determine whether TCF4 restoration could rescue these deficits, we assessed locomotor activity and exploratory behavior in R342X and C99F-m mice using the open-field test (**Fig. 6**). Both mutant lines showed increased locomotor activity compared with WT controls. R342X mice traveled significantly greater distances than WT mice (WT: ∼45 m, R342X: ∼60 m), while C99F-m mice also showed increased distance traveled (WT: ∼40 m, C99F-m: ∼50 m). Both BFLS mouse models spent significantly more time in the center of the arena than WT controls (R342X mice ∼40 s, C99F-m mice ∼30 s, and ∼25 s in WT) and significantly less time in the corners than WT controls. Together, these findings demonstrate altered locomotor activity in both BFLS models.

**Figure 6:**
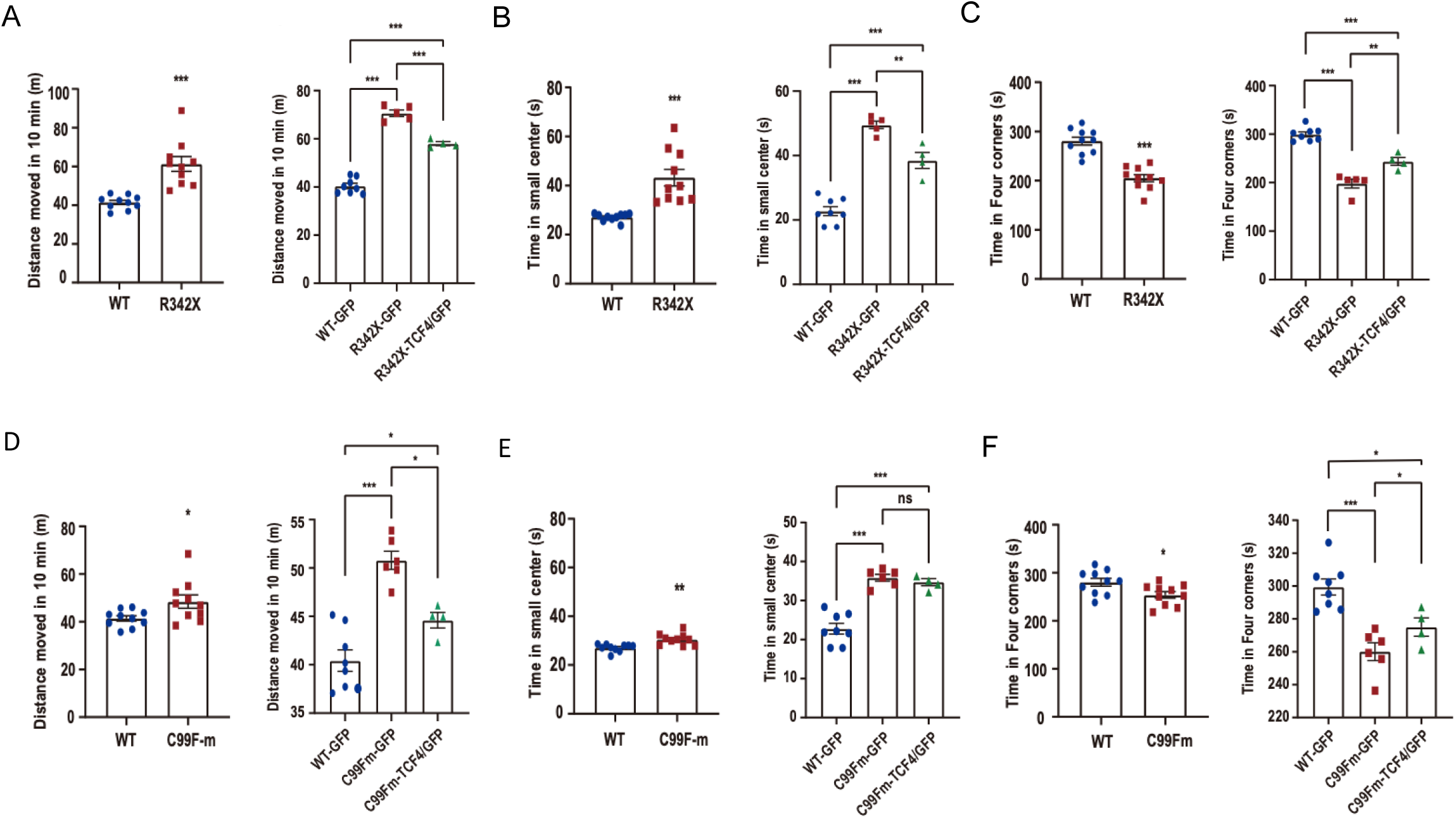
TCF4 restoration rescues altered locomotor activity in BFLS mouse models. (A) Distance traveled in the open-field test for R342X and WT mice. (B) Distance traveled in R342X mice expressing GFP or TCF4; TCF4 expression normalized locomotor activity toward WT levels. (C) Time spent in the center zone and four corners for R342X mice, showing partial restoration of exploratory behavior with TCF4 expression. (D) Distance traveled in the open-field test for C99F-m and WT mice. (E) Distance traveled in C99F-m mice expressing GFP or TCF4; TCF4 expression significantly improved locomotor activity. (F) Time spent in the center zone and four corners for C99F-m mice, showing restoration of exploratory behavior with TCF4 expression. Data are presented as mean ± SEM. Statistical analysis was performed using one-way ANOVA with a post-hoc test. *p < 0.05, **p < 0.01, ***p < 0.001.

We next tested whether restoration of TCF4 could rescue these behavioral abnormalities. In R342X mice, TCF4 expression significantly improved the altered locomotor phenotype. R342X- TCF4/GFP mice showed reduced distance traveled toward WT levels (∼45 m compared with ∼60 m in R342X-GFP; **Fig. 6B**). TCF4 expression also partially normalized center exploration and increased the time spent in the corners compared with R342X-GFP (**Fig. 6C**). Similarly, C99F-m- TCF4/GFP mice showed significant improvement in distance traveled, center exploration, and corner exploration compared with C99F-m-GFP (**Fig. 6D-F**).

### TCF4 restoration improves impaired object exploration in BFLS

We next assessed object exploration using the novel object recognition paradigm (**Fig. 7**). Both R342X and C99F-m mutant mice exhibited reduced interaction with familiar objects compared with WT controls. During the novel object trial, both mutant lines also showed significantly reduced interaction with the novel object compared with WT controls. These findings demonstrate impaired object exploration in both BFLS mouse models.

**Figure 7:**
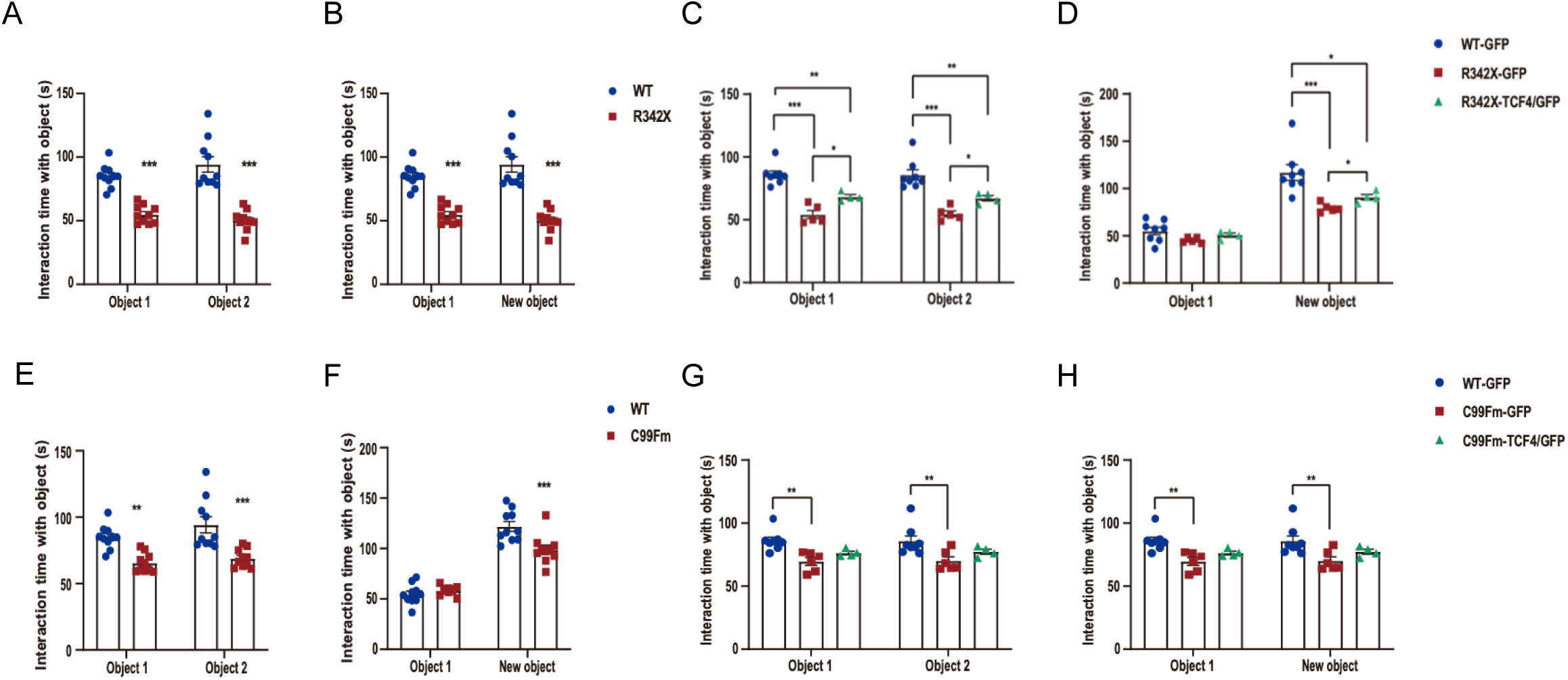
TCF4 restoration rescues impaired object exploration in R342X and C99F-m BFLS mice. (A) Interaction time with Object 1 and Object 2 in R342X and WT mice; R342X mice showed significantly reduced object exploration. (B) Interaction time with Object 1 and a novel object in R342X and WT mice; R342X mice exhibited reduced novel object interaction. (C) Interaction with familiar objects (Object 1 and Object 2) in R342X mice expressing GFP or TCF4; TCF4 expression significantly increased object interaction. (D) Interaction with familiar and novel objects in R342X mice; TCF4 expression substantially improved novel object exploration. (E) Interaction time with familiar and novel objects in C99F-m and WT mice; C99F-m mice showed reduced object exploration. (F) Interaction with Object 1 and novel object in C99F-m and WT mice. (G) Interaction with familiar objects in C99F-m mice expressing GFP or TCF4; TCF4 expression increased exploration. (H) Interaction with familiar and novel objects in C99F-m mice; TCF4 expression significantly improved novel object interaction. Data are presented as mean ± SEM. Statistical analysis was performed using one-way ANOVA with a post-hoc test. *p < 0.05, **p < 0.01, ***p < 0.001

We next asked whether TCF4 restoration could improve these behavioral abnormalities. R342X-TCF4/GFP mice showed significantly increased interaction with both familiar and novel objects compared with R342X-GFP controls (**Fig. 7C, D**). Similarly, C99F-m-TCF4/GFP mice showed increased interaction with familiar and novel objects compared with C99Fm-GFP (**Fig. 7G, H**). Together, these findings demonstrate that TCF4 restoration during cortical development improves altered exploratory and object-directed behaviors in both BFLS mouse models.

### PHF6 and TCF4 correlation in human brain development

Our analysis on aNSC reveals that PHF6-dependent regulation of TCF4 has consequences extending beyond the embryonic stage, suggesting that such mutation-induced perturbations propagate across the lifespan and may contribute to persistent neurological dysfunction in the adult brain. To assess whether this relationship is conserved in humans, we analyzed bulk RNA-seq profiles from the human ventral frontal cortex across developmental time points^56^. Consistent with the findings from the mouse dataset, *PHF6* and *TCF4* exhibited parallel expression trajectories throughout human cortical development (**Fig. 8A, B**).

**Figure 8:**
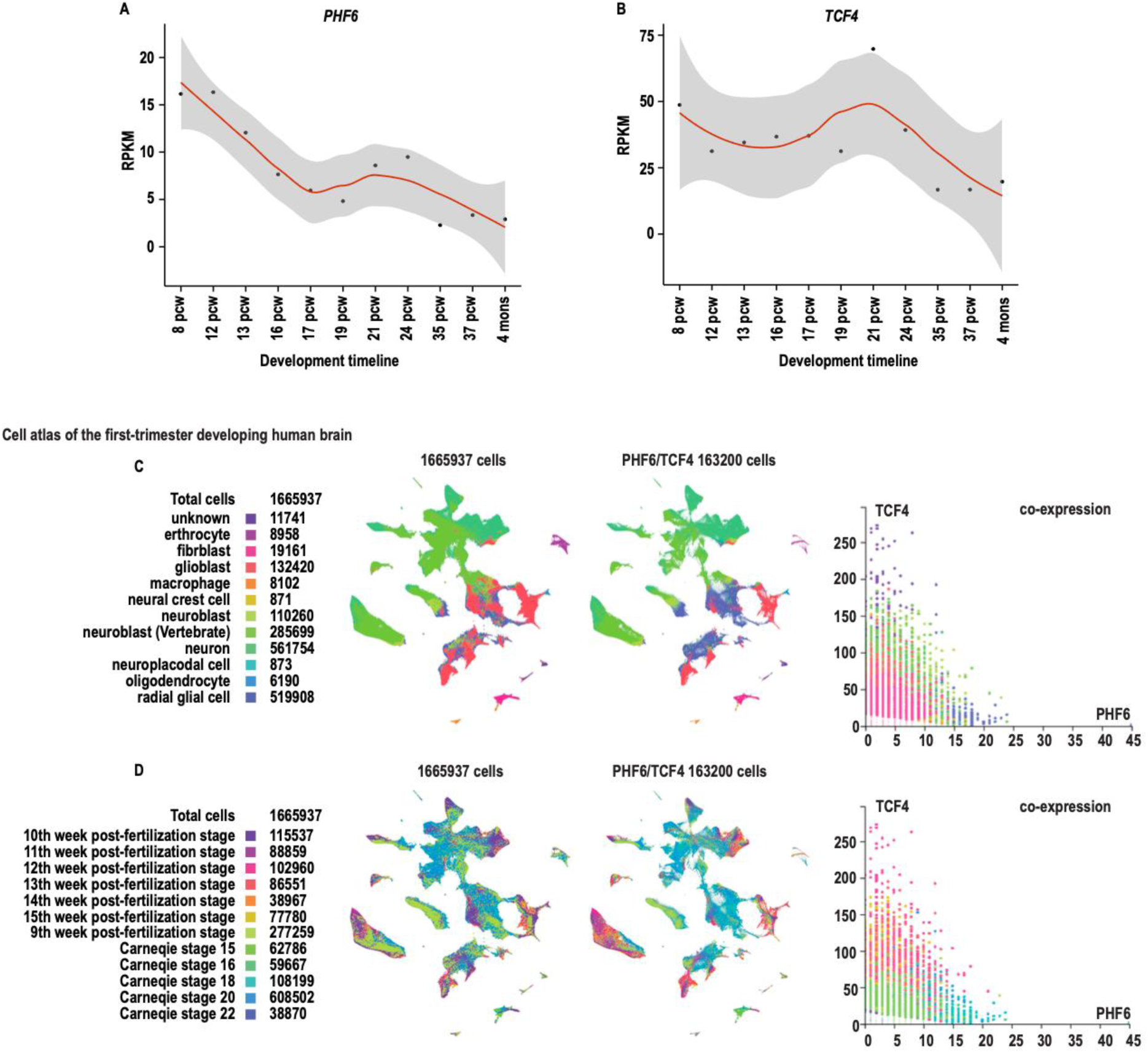
Analysis of *PHF6* and *TCF4* expression across development in human. (A-B) Analysis of *PHF6* and *TCF4* expression in the human cortex. Plots show average reads per kilobase per million mapped reads (RPKM) across human developmental time (post-conceptual weeks; pcw) for (A) *PHF6* and (B) *TCF4*. Expression values were obtained from publicly available RNA-seq data generated from the human ventral frontal cortex (VFC) as part of the Allen Brain Atlas BrainSpan dataset56. (C) UMAP embedding of 1,665,937 cells from the first-trimester human brain, colored by annotated cell types. Scatter plot of PHF6 versus TCF4 expression across all cells, illustrating their co-expression relationship. (D) UMAP embedding of all cells colored by developmental stage, including post-fertilization weeks (9–15) and Carnegie stages (15–22). Cell counts for each stage are provided. Scatter plot of PHF6 versus TCF4 expression stratified by developmental stage, demonstrating stage-specific patterns of co-expression. Data obtained from CELLxGENE datasets ^56^.

We further analyzed the co-expression of *PHF6* and *TCF4* in human brain development to confirm our observation in PHF6 and BFLS mouse models. We employed a large-scale snRNA-seq atlas of the first-trimester human brain comprising 1,665,937 cells, accessed through the CELLxGENE platform. UMAP visualization revealed a diverse cellular composition, including radial glial cells (519,908 cells), neurons (561,754 cells), neuroblasts (285,699 cells), glioblasts (132,420 cells), and fibroblasts (19,161 cells) (**Fig. 8C**). Within this dataset, 163,200 cells were identified to express both *PHF6* and *TCF4*. These cells were distributed across multiple cell types and preferentially localized to distinct clusters within the UMAP embedding, indicating that co-expression is not restricted to a single lineage but is broadly present across progenitor and neuronal populations during early human brain development (**Fig. 8C**).

Stratification of the dataset by developmental stage, spanning Carnegie stages 15–22 and post-fertilization weeks 9 to 15, demonstrated that *PHF6*/*TCF4* co-expressing cells were present across all stages examined, with the largest cell numbers contributed by Carnegie stage 20 (608,502 cells) and the 10th week post-fertilization stage (115,537 cells) (**Fig. 8D**). Scatter plot analysis of *PHF6* versus *TCF4* expression across co-expressing cells revealed a consistent positive co-expression relationship maintained throughout these developmental windows, supporting a coordinated regulatory relationship between the two factors during human corticogenesis. Collectively, these findings suggest that the *PHF6*-*TCF4* signalling axis may operate as a developmentally conserved transcriptional mechanism in the regulation of cell fate.

## Discussion

In this study, we identify a previously uncharacterized PHF6–*Tcf4* transcriptional mechanism that governs NSC fate. Using complementary BFLS and PHF6 mouse models, including R342X, C99F-m and a *Phf6* cKO, we demonstrate that PHF6 directly binds the *Tcf4* promoter to drive its expression in eNSCs, and that loss of PHF6 function consistently suppresses TCF4 levels across all models. *Tcf4* knockdown phenocopies PHF6 deficiency by enhancing NSC self-renewal and stemness marker expression, while forced TCF4 expression rescues these defects, establishing TCF4 as a bona fide functional effector downstream of PHF6. These findings provide the first evidence linking TCF4 dysregulation to BFLS pathogenesis.

PHF6 and TCF4 have each been independently studied in neurodevelopment, yet their functional relationship had not previously been explored. PHF6 operates through interactions with the PAF1 complex, UBF1, and NuRD complex to regulate neuronal migration and chromatin organization^12, 22, 46–49^, while TCF4 plays critical roles in neural progenitor proliferation, differentiation, and migration, exhibiting layer-specific expression in the developing neocortex and high expression in adult neurogenic niches^24, 28–30^. Despite their distinct molecular mechanisms, PHF6 as a genetic and epigenetic modifier, and TCF4, as a transcription factor, appear to operate on the same signalling pathway in the regulation of stem cells. Our single-cell RNA-seq correlation analyses reveal positive co-expression of *Phf6* and *Tcf4* across progenitor populations throughout cortical development, consistent with their shared peak expression during mid-to-late embryogenesis. The data presented here establish TCF4 as a critical transcriptional effector in the PHF6-dependent program governing NSC fate.

Importantly, our findings demonstrate that dysregulation of the PHF6-*Tcf4* pathway translates into significant behavioral and cognitive abnormalities in vivo. Comprehensive behavioral characterization of BFLS mouse models^39, 51^ revealed hyperlocomotor activity and marked impairments in object exploration and novelty recognition. Critically, IUE-mediated restoration of TCF4 expression during cortical development rescued these behavioral and cognitive abnormalities, normalizing locomotor activity while improving object exploration and novelty recognition in both R342X and C99F-m BFLS mice.

Cortical neurogenesis requires tightly regulated transitions from NSCs through intermediate progenitors to neurons^50, 51^. Similar to PHF6, TCF4 restricts NSCs self-renewal, suggesting that PHF6 deficiency disrupts NSC transitions via downregulation of TCF4, which is likely expected to impair progenitor commitment, and consequently alter neuronal output and induce cortical organization defects associated with BFLS. At the circuit level, disrupted cortical neurogenesis and consequent alterations in neural circuit development may contribute to the cognitive and behavioral deficits observed in BFLS models. Notably, the dissociation between hyperlocomotor activity and reduced exploratory behavior suggests that these abnormalities may reflect preferential disruption of neural circuits involved in attention, cognitive engagement, and executive function.

These findings align with reports that *Tcf4* disruption causes abnormal cortical development^26^ and that TCF4 loss-of-function in human cerebral organoids alters progenitor dynamics^44^. However, it remained to be determined whether TCF4 restoration can ameliorate behavioral phenotype and defects in brain connectivity in BFLS. Similarly, this study raises future questions on whether BFLS patients may exhibit psychiatric disorders associated with TCF4 deregulation.

A particularly important implication of these findings relates to the broader landscape of TCF4-associated disorders. Heterozygous loss-of-function mutations in *TCF4* cause Pitt-Hopkins syndrome^30–32^, and *TCF4* polymorphisms are robustly associated with schizophrenia and autism spectrum disorder by genome-wide association studies^34, 35, 52^. Our findings open a new avenue of research to establish and delve deeper into shared molecular mechanisms across these disorders and, more specifically, linking BFLS to a wider spectrum of neurodevelopmental and psychiatric disorders.

## Materials and methods

### Animals

All animal experiments received approval from the Animal Care Committee (ACC) at the University of Ottawa, Ottawa, Ontario, Canada. Mice were housed under standard conditions in a pathogen-free facility with ad libitum access to food and water.

### Mouse Models

The R342X mouse line was created via CRISPR/Cas9 technology, producing a truncated PHF6 protein^9–12, 17, 18, 40^. This line was maintained by crossing R342X female heterozygous (HET) mice with C57BL/6J wild-type (B6 WT) male mice. Hemizygous (HEMI) males served as experimental subjects, while B6 WT males were used as controls.

The C99F-m mouse model was generated using CRISPR/Cas9, where cysteine at position 99 is substituted with phenylalanine (C99F) through a nucleotide change at at nt.29 G > T^53^. This line was propagated by breeding C99F-m female HET mice with B6 WT male mice. HEMI males were designated as experimental animals, with B6 WT males as controls.

The brain-specific *Phf6* knockout (*Phf6^-/Y^/Nestin-CreERT2^+^*) strain was generated by crossing Phf6^fl/fl^ female mice with Nestin-CreERT2^+^ male mice^54^. Cre recombinase activity was induced through oral administration of Tamoxifen (Sigma-Aldrich, T5648) to pregnant dams at E12, with embryo collection occurring 48 hours post-treatment. *Phf6^-/Y^/Nestin-CreERT2^+^*mice were analyzed alongside *Phf6^loxP/Y^/Nestin-CreERT2^−^* control mice that also received tamoxifen treatment. Male mice were utilized throughout these experiments.

### Genotyping Procedures

Mouse tissue samples (ear biopsies) were lysed in alkaline lysis buffer (25 mM NaOH, 0.2 mM EDTA, pH 12) and incubated at 95°C for 30 minutes. Samples were subsequently neutralized with an equivalent volume of neutralization buffer (40 mM Tris-HCl, pH 5.0).

For C99F-m and R342X genotyping, PCR reactions (25 μL total volume) contained: 2.5 μL 10× PCR buffer, 0.2 μL 25 mM dNTP, 6.5 μL betaine, 1 μL each of 10 μM forward and reverse/mutation primers, 0.2 μL Klentaq Thermostable DNA Polymerase (Jena Bioscience, #PCR-217L), 12.6 μL RNase-free H₂O, and 1 μL DNA template.

For Phf6^fl/fl^ genotyping, PCR reactions (15 μL total) were prepared with: 7.5 μL 2× Green PCR Master Mix (ZmTech Scientific, #S2100G), 0.4 μL each of 10 μM forward and reverse primers, 5.7 μL RNase-free H₂O, and 1 μL DNA template.

Nestin-CreERT2 genotyping reactions (15.5 μL total) consisted of: 7.5 μL 2× Green PCR Master Mix (ZmTech Scientific, #S2100G), 1.5 μL each of 0.5 μM primers (oIMR1084, oIMR1085, oIMR7338, and oIMR7339), 0.975 μL 6.5% glycerol, and 1 μL DNA template.

PCR amplification was performed in a Bio-Rad T100 Thermal Cycler. For C99F-m, R342X, and *Phf6^Loxp/Loxp^*, the cycling conditions were: initial denaturation at 95°C for 2 minutes; 34 cycles of 95°C for 30 seconds, 60°C for 30 seconds, and 72°C for 30 seconds; followed by final extension at 72°C for 4 minutes.

Nestin-CreERT2 samples underwent: 94°C for 2 minutes; 10 touchdown cycles of 94°C for 20 seconds, 65°C for 15 seconds (decreasing 0.5°C per cycle), and 68°C for 10 seconds; 28 cycles of 94°C for 15 seconds, 60°C for 15 seconds, and 72°C for 10 seconds; with final extension at 72°C for 2 minutes.

Nestin-Cre genotyping used: 94°C for 2 minutes; 35 cycles of 94°C for 20 seconds, 60°C for 20 seconds, and 72°C for 25 seconds; followed by 72°C for 2 minutes.

Amplified products were analyzed by electrophoresis on 3% agarose gels at 100 V for 40 minutes (C99F-m, R342X, Nestin-CreERT2, and Nestin-Cre) or 60 minutes (*Phf6^fl/fl^*).

### Tamoxifen Induction

For temporal control of Cre recombinase in Phf6^fl/fl^/Nestin-CreERT2 mice, pregnant dams at gestational day E12 received a single 0.1 mL oral gavage of Tamoxifen (Sigma-Aldrich, T5648) at 20 mg/mL concentration using a 1 mL syringe fitted with a 22-gauge feeding needle (Instech Solomon, #FTP-22-25-5).

### Embryonic neural stem cell culture

Embryonic neural stem cells (eNSCs) were obtained by culturing whole brains isolated from E14 mice (excluding cerebellum). Pregnant mice were euthanized, and embryos were harvested by dissecting the uterine horns. Embryos were transferred into cold 1× Hank’s Balanced Salt Solution (HBSS). Brain tissues were dissected from embryos and immediately placed into a 15mL Falcon containing 1mL cold 1× HBSS. The tissue was allowed to settle, HBSS was replaced with 1 mL fresh HBSS for washing, then replaced again with 1 mL of stem cell medium (SCM) containing 1:1 DMEM-F12 (Wisent, 319-005-CL) (ThermoFisher, 31765035), 50 units/mL penicillin-streptomycin (Wisent, 450-201-EL), 1× B-27 supplement (Invitrogen, 17504044), 10 mg/mL Heparin (Stemcell Technologies, 07980), 100µg/mL EGF (Cell Signaling, 5331SC), 25µg/mL bFGF (Abbiotec, 600182). The tissue was triturated 10 times with a P1000 pipette, then 10 times with a P200 pipette. The single cell was plated in 6 mL of SCM and incubated for 6–7 days, allowing neurospheres to grow.

### Adult neural stem cell culture

Adult neural stem cells (aNSCs) were cultured as we established before^42^. Briefly, aNSCs were extracted from mouse brains, the subventricular zone (SVZ) and subgranular zone (SGZ). The tissue was dissected and transferred to sterile microcentrifuge tubes. Papain working solution (PWS, 15µL of Papain suspension enzyme is added to 1 mL of 0.53 mM EDTA and 10µL L-Cysteine) was filter-sterilized using a 0.22 µm syringe filter into the Falcon. Dissected tissue was transferred into Falcon using a plastic pipette pre-wetted with PWS. Tissue was mechanically dissociated by pipetting until freely entering the tip, followed by 20 rounds of trituration using a P1000 pipette set to 800 µL. Samples were incubated at 37 °C for 10 minutes and homogenized again (20 cycles, P1000 set to 800 µL). DNase I (5 µL of 10 ng/µL) was added to each sample, mixed by gentle flicking, and incubated at 37 °C for 20 minutes. Samples were homogenized again (20 times), followed by a second DNase treatment using the same conditions, flicked gently, and incubated at 37 °C for 15 minutes. This was followed by two additional cycles of 15-minute incubations and 20 homogenization strokes. After the final incubation, 400 µL of trypsin inhibitor (1 mg/mL) was added to each sample to terminate enzymatic digestion. Five millilitres of base media containing 3:1 of DMEM-F12 and 250 µL of 1% penstrep were added to each tube. For each sample, the cell suspension was filtered through a 40 µm cell strainer pre-wetted with 8 mL of base media placed on a fresh 50 mL Falcon tube; the 8 mL was discarded before slowly pouring the sample through the strainer and centrifuged at 1000 rpm for 10 minutes. The supernatant was carefully aspirated, and the cell pellet was resuspended in 1 mL of proliferation media containing 3:1 of DMEM-F12, 500 µL of 1% penstrep, 1mL 2% B27, and 20µL of 50µg/mL mouse EGF. Cells were resuspended and transferred to a T75 flask containing 29mL of fresh proliferation media and incubated for 14 days, allowing neurospheres to grow.

### Extreme Limiting Dilution Assay (ELDA) and Limiting Dilution Assay (LDA)

Extreme Limiting Dilution Assay (ELDA) and Limiting Dilution Assay (LDA) were performed as described before^36, 42^. Briefly, single-cell NSCs suspensions were prepared using Accumax. Single cells were counted and plated at varying densities in 96-well plates. For ELDA, sphere formation was counted 7 days post-plating, and stem cell frequency (SCF) was calculated. For LDA, spheres were counted 7 days post-plating, in triplicate. ELDA data was analyzed through http://bioinf.wehi.edu.au/software/elda/^33 43^.

### Quantitative real-time PCR

RNA was isolated from cells and brain tissue with Trizol (Invitrogen) according to the manufacturer’s instructions. Reverse transcription of RNA was performed using 5x All-In-One RT MasterMix cDNA synthesis (Abm, G492). Quantitative real-time PCR was performed using SsoAdvanced Universal SYBR®Green Supermix (Bio-Rad, 1725271). Samples were incubated at 25 °C for 10 min, followed by incubation at 42 °C for 15 min, and finally at 85 °C for 5 min to inactivate the reaction.

### Immunoblotting

Protein lysates were prepared from brain tissue using RIPA lysis buffer containing protease and phosphatase inhibitors (ThermoFisher Scientific, A32959). The concentration of proteins was analyzed by the Bradford Assay (Bio-Rad) with BSA standard. PVDF membranes were activated in methanol for 5 min and then blocked in 5% BSA in TBST. Membranes were probed with anti- PHF6 (NOVUS, NB100-68262, 1:1000), anti-TCF4 (ThermoFisher, PA5-75987, 1:1000), anti-SOX2 (Abcam, ab97959, 1:250), anti-NESTIN (Santa Cruz, sc-23927, 1:100) or (R&D Systems, MAB2736, 1:500), anti-beta-Actin (Sigma-Aldrich, a5316, 1:2000), alpha-Tubulin (Abcam, 9074, 1:5000), overnight at 4 °C, followed by HRP-conjugated secondary antibody, anti-rabbit IgG HRP (Bio-Rad, 1706515) or anti-mouse IgG HRP (Bio-Rad, 1706516) for 2 h at room temperature. Proteins were visualized using ECL (Bio-Rad), and images were captured with a Chemidoc imaging system (Bio-Rad).

### Chromatin immunoprecipitation (ChIP)

Cell pellets were collected and washed with PBS supplemented with protease inhibitors (Thermo Fisher Scientific, #A32959) before fixation. Cross-linking was performed using 1% formaldehyde in PBS for 10 minutes, followed by quenching with 0.125 M glycine in PBS for 5 minutes at room temperature. Washing, fixation, and quenching were done in 15 mL tubes while rotating at RT. After quenching, cells were washed twice with PBS containing protease inhibitors. Cell pellets were collected by spinning at 150 × *g* for 5 min at 4 °C. Next, cell pellets were dissolved in ChIP lysis buffer (40 mM Tris-HCl, pH 8.0, 1.0% Triton X-100, 4 mM EDTA, 300 mM NaCl) containing protease inhibitors. Chromatin fragmentation was performed through tip sonication on ice. Cell lysates were centrifuged at 12,000 × g for 15 minutes at 4°C, and the resulting supernatant was diluted 1:1 with ChIP dilution buffer (40 mM Tris-HCl, pH 8.0, 4 mM EDTA, protease inhibitors).

Immunoprecipitation (IP) was done using a PHF6 antibody (Novus Biological, NB100-68262). Antibody-protein-DNA complexes were collected, washed, and then eluted. Reverse cross-linking was done as established before ^43^. Immunoprecipitated DNA was quantified by qPCR, and binding enrichment was calculated as a percentage relative to input DNA.

### Dual-luciferase reporter assay

The PHF6 binding regions (based on ChIP-seq peaks) were cloned into the pGL4.23 (Promega) vector to generate the *TCF4* luciferase reporter genes by digesting the plasmid and the annealed primer pair using KpnI (NEB, #R3142) and HindIII (NEB, #R3104) then ligating them with T4 DNA ligase (NEB, #M0202L). The constructs were confirmed by DNA sequencing. eNSCs were electroplated with the TCF4-pGL4.23 or the empty pGL4.23. Luciferase assays were performed 48h after electroporation with the Dual-Luciferase Reporter Assay system (Promega, #E1910) with Synergy H1 BioTek (luminance mode). In all experiments, cells were electroplated with a Renilla firefly reporter control, and the firefly luminescence signal was normalized to the Renilla luminescence signal.

### Immunofluorescence staining of tissue

Immunofluorescence was performed as before^36^. After blocking, sections were subjected to the following primary antibody for TCF4 immunofluorescence (overnight at 4 °C): anti-TCF4 (1:100, ThermoFisher, PA5-75987), and DAPI (1:1000, ThermoFisher, D1306). Secondary antibodies were applied for 45 min at room temperature in a humid chamber.

### siRNA

*Tcf4* was knocked down using short interfering RNA (siRNA) in eNSCs and aNSCs with ON TARGET-plus SMART pool mouse *Tcf4* siRNA (Santa Cruz, sc-61658) and ON TARGET-plus non-targeting pool (Santa Cruz, #sc-36869) at a concentration of 100 nM. siRNA was nucleofected into NSCs (10⁶ cells) and cultured in specific media at 37 °C in a humidified atmosphere of 5% CO₂.

### Plasmid

A lentiviral vector for overexpressing the TCF4 transcription factor open reading frame (ORF), tagged with a unique 24-bp barcode to enable identification in pooled screens, was obtained from Addgene (plasmid #141560).

### In utero electroporation

In utero electroporation (IUE) was performed at E14.5 to induce TCF4 overexpression in the developing mouse brain, as previously described^57^. Pregnant dams were anesthetized, and the uterine horns were gently exteriorized to expose the embryos. A TCF4 expression plasmid mixed with a fluorescent tracer dye was injected into the lateral ventricle of each embryo. Successful injection was confirmed by visualization of the dye within the ventricular space.

Following DNA injection, the embryo was subjected to in utero electroporation using paddle electrodes positioned across the uterus. Electrical pulses were applied to facilitate plasmid uptake by neural progenitor cells in the developing brain. The uterine horns were subsequently returned to the abdominal cavity, and the abdominal and skin incisions were closed. The dams were allowed to recover under standard postoperative care. TCF4-electroporated embryos were allowed to develop to the desired postnatal age for subsequent behavioral and histological analyses. The fluorescent tracer was used to identify and confirm the targeted/electroporated regions.

### Behavioral testing

All behavioral assessments were performed in the Behavior Core Facility at the University of Ottawa following established guidelines. Before testing, mice were allowed to habituate to the testing environment for 1 h and were then returned to their home cage. All testing equipment was thoroughly cleaned with 70% ethanol between animals.

### Open field

Open field testing was performed to assess anxiety-related behavior in wild-type (C57BL/6), R342X, and C99F-m mice. Mice were individually placed in the center of a 45 × 45 × 45-cm open-field chamber and allowed to freely explore for 10 min under illumination of approximately 300 lux. Behavioral activity was recorded and analyzed using the EthoVision XT video tracking system. During the 10-min test period, total distance traveled, time spent in the four corner regions, and time spent in the central zone were quantified. Ten mice per genotype were tested. The arenas were cleaned between animals to minimize the influence of odors from previously tested mice.

### Novel objective recognition

Novel object recognition testing was performed to assess recognition memory using two identical testing chambers. Mice were habituated to the testing room for 1 h before each testing session. Behavioral activity was recorded using the EthoVision XT video tracking system under red-light illumination. On the first day, mice were first individually placed in an empty testing chamber and allowed to habituate for 5 min. Following habituation, mice were returned briefly to their home cages. Mice were then placed individually in the chamber containing two identical objects, cups or funnels, and allowed to freely explore for 10 min. Following this familiarization trial, mice were returned to their home cages.

On the second day, mice were again habituated to the testing room for 1 h before testing. Each mouse was then placed in the chamber containing one familiar object and one novel object (a cup and a funnel). The mice were allowed to freely explore the chamber for 10 min, during which interactions with both objects were recorded using EthoVision XT. The time spent interacting with the novel and familiar objects was quantified, and preference for the novel object was used as a measure of recognition memory.

### Leveraging published RNA sequencing datasets

Single cell RNA-seq data from the developing mouse cerebral cortex were obtained from Di Bella et al (2021)^37^. Log-normalized counts, cell type annotations and UMAP coordinates were retrieved from the original publication and used to generate UMAP plots. For the correlation analyses, MAGIC^55^ (v.2.0.3) was applied to obtain imputed gene-expression values, and Pearson correlation coefficients were computed using the imputed matrix. For human bulk RNA-seq profiles, normalized RPKM (Reads per Kilobase Million) values were retrieved from the Allen Brain Atlas BrainSpan dataset^56^ and filtered to include only samples from the ventral frontal cortex (VFC). Average RPKM values were calculated for each developmental time point. All analyses and plots were generated using R (v.4.3.1).

### Analysis of PHF6 and TCF4 co-expression in the developing human brain using CELLxGENE

To investigate the temporal co-expression of *PHF6* and *TCF4* during human brain development, we accessed publicly available single-nucleus RNA sequencing (snRNA-seq) datasets through the CELLxGENE platform (https://cellxgene.cziscience.com). For the analysis of the first-trimester developing human brain, a dataset comprising 1,665,937 cells was retrieved. Cell type annotations and developmental stage classifications were as provided in the original dataset metadata available through CELLxGENE. Dimensionality reduction and visualization were performed using Uniform Manifold Approximation and Projection (UMAP). Cells were visualized and colored by annotated cell type or by developmental stage to examine the cellular composition and temporal distribution across the datasets.

To assess the co-expression relationship between *PHF6* and *TCF4*, cells simultaneously expressing both genes were identified and extracted from each dataset. For the human first-trimester brain dataset, 163,200 double-positive cells (PHF6^+^/TCF4^+^) were identified. These co-expressing cells were projected onto UMAP embeddings for spatial visualization. Scatter plots of *PHF6* versus *TCF4* normalized expression values were generated for the double-positive subsets and further stratified by developmental stage and cell type to resolve stage-specific and lineage-specific co-expression patterns.

### Statistical analysis

Statistical analysis was performed with software Prism 8. Two-tailed unpaired Student t-tests were used to compare two conditions. One-way ANOVA was used to analyze multiple groups. Data are shown as mean with standard error of mean (mean ± SEM). *p*-values of less than 0.05 were considered significant and were marked with one asterisk (*). *p*-values of less than 0.01 are denoted by **, and *p-values* of less than 0.001 are denoted by ***. All data presented are from 3 or more independent biological (n) replicates (*n* ≥ 3). Methods of statistical analysis and *p*-values employed are reported in the corresponding figure legends.

### Graphics

The graphics were created with BioRender.com.

## Acknowledgments

We are grateful to all staff from the Animal Care and Veterinary Service (ACVS) at the University of Ottawa for their continuous support. We also thank the University of Ottawa Behavioural Core Facility for providing access to behavioural testing equipment and technical support.

